# A single dietary factor, daily consumption of a fermented beverage, can modulate the gut microbiome within the same ethnic community

**DOI:** 10.1101/2023.01.18.524612

**Authors:** Santanu Das, Ezgi Özkurt, Tulsi Kumari Joishy, Dibyayan Deb, Ashis K. Mukherjee, Falk Hildebrand, Mojibur R. Khan

**Author notes:** Both the authors have contributed equally (Santanu Das, Ezgi Özkurt). Corresponding authors: Dr. M. R. Khan, Life Sciences Division, Institute of Advanced Study in Science and Technology (IASST), Guwahati-781035, Assam, India., Dr. Falk Hildebrand, Gut Microbes & Health, Quadram Institute Bioscience, Norwich Research Park, Norwich, Norfolk, NR4 7UA, UK & Earlham Institute, Norwich ResearchPark, Norwich, Norfolk, NR4 7UA, UK.

## Abstract

In this study, the impact of traditional rice-based fermented alcoholic beverages (*Apong*) on the gut microbiome and health of the *Mishing* community in India was examined. Two groups that consumed one of these beverages were compared to a control group that did not consume either beverage. Gut microbial composition was analyzed by sequencing 16S rRNA of fecal metagenomes and analyzing untargeted fecal metabolites, and short-chain fatty acids (SCFAs). We also collected data on anthropometric measures and serum biochemical markers. Our results showed that *Apong* drinkers had higher blood pressure, but lower blood glucose and total protein levels than other non-drinkers. Also, gut microbiome composition was found to be affected by the choice of *Apong*, with *Apong* drinkers having a more diverse and distinct microbiome compared to non-drinkers. *Apong* drink type or being a non-drinker explained even a higher variation of fecal metabolome composition than microbiome composition and *Apong* drinkers had lower levels of the SCFA isovaleric acid than non-drinkers. Overall, this study shows that a single dietary factor can significantly impact the gut microbiome of a community and highlights the potential role of traditional fermented beverages in maintaining gut health.

## Introduction

The gut microbiome is a collection of microorganisms in the human intestine that performs many important functions, including digestion, transformation of nutrients, and immune system regulation^1^. A well-balanced gut microbiome composition can provide health benefits, while imbalances can lead to disorders related to metabolism and immunity^2^. The gut microbiome is influenced by various factors, including diet, medication, and age. Diet is a particularly important factor that affects the gut microbiome composition and its interactions with the host^5–7^. Fermented foods and beverages, such as yogurt, kefir, fermented cottage cheese, kimchi, and kombucha tea, are rich in microorganisms that can have an effect on the gut microbiome and increase overall microbial diversity^8–12^.

Rice-based fermented beverages are an important part of the diet and cultural heritage of Mongoloid communities in South-East Asia^13^. In Assam, India, the Mishing community consumes two types of rice-based alcoholic beverages called *Poro Apong* and *Nogin Apong. Poro Apong* is prepared with roasted rice, ash of rice husk, and a starter cake, and undergoes solid-state fermentation for 7-10 days. It is filtered through ash to produce a dark, clear liquid with physical and sensory properties similar to stout beer. *Nogin Apong* is made with steamed rice, and has physical and sensory properties similar to *Maakoli*, a fermented beverage from South Korea. Within the Mishing community, some subgroups consume only one type of *Apong*, despite having similar lifestyles and dietary habits. We previously found that *Nogin Apong* had a diverse array of lactic acid bacteria and was rich in saccharides and amino acids, while *Poro Apong* was dominated by *Lactobacillus*^14^. In this study, we aimed to determine how the different compositions of microbes and metabolites in these two types of *Apong* may affect the gut microbiome in volunteers from the same ethnicity who have similar dietary patterns, but differ in their choice of *Apong*.

## Results

### *Apong* drinkers have distinct levels of biochemical markers compared to non-drinkers, including differences in blood pressure, glucose, protein levels, and liver enzymes

Most individuals in this study had normal physiological and biochemical test results regardless of their lifestyles and dietary habits. However, blood pressure was significantly higher in *Poro* drinkers compared to the controls. **(Figure 1B, a-b)**. Both *Nogin* and *Poro* drinkers had lower total protein and albumin levels in their blood than non-drinkers **(Figure 1B, c-d)**, with *Nogin* drinkers having a lower albumin to globulin ratio **(Figure 1B, f)**. Both *Nogin* and *Poro* drinkers also had lower blood glucose levels than non-drinkers **(Figure 1B, g)**, which is consistent with the blood glucose-lowering properties of fermented foods and beverages^15^. Lipid levels, as measured by triglycerides, were within the normal range and comparable in all participants **(Figure 1B, h)**. *Poro* drinkers had significantly lower levels of SGPT **(Figure 1B, i)** and SGOT **(Figure 1B, j)**, which is a sign of a healthy liver^16^, and lower levels of bilirubin total and bilirubin direct **(Figure 1B, k-l)**. Although high bilirubin levels can indicate liver damage, low levels are not a concern for health.

**Figure 1.**
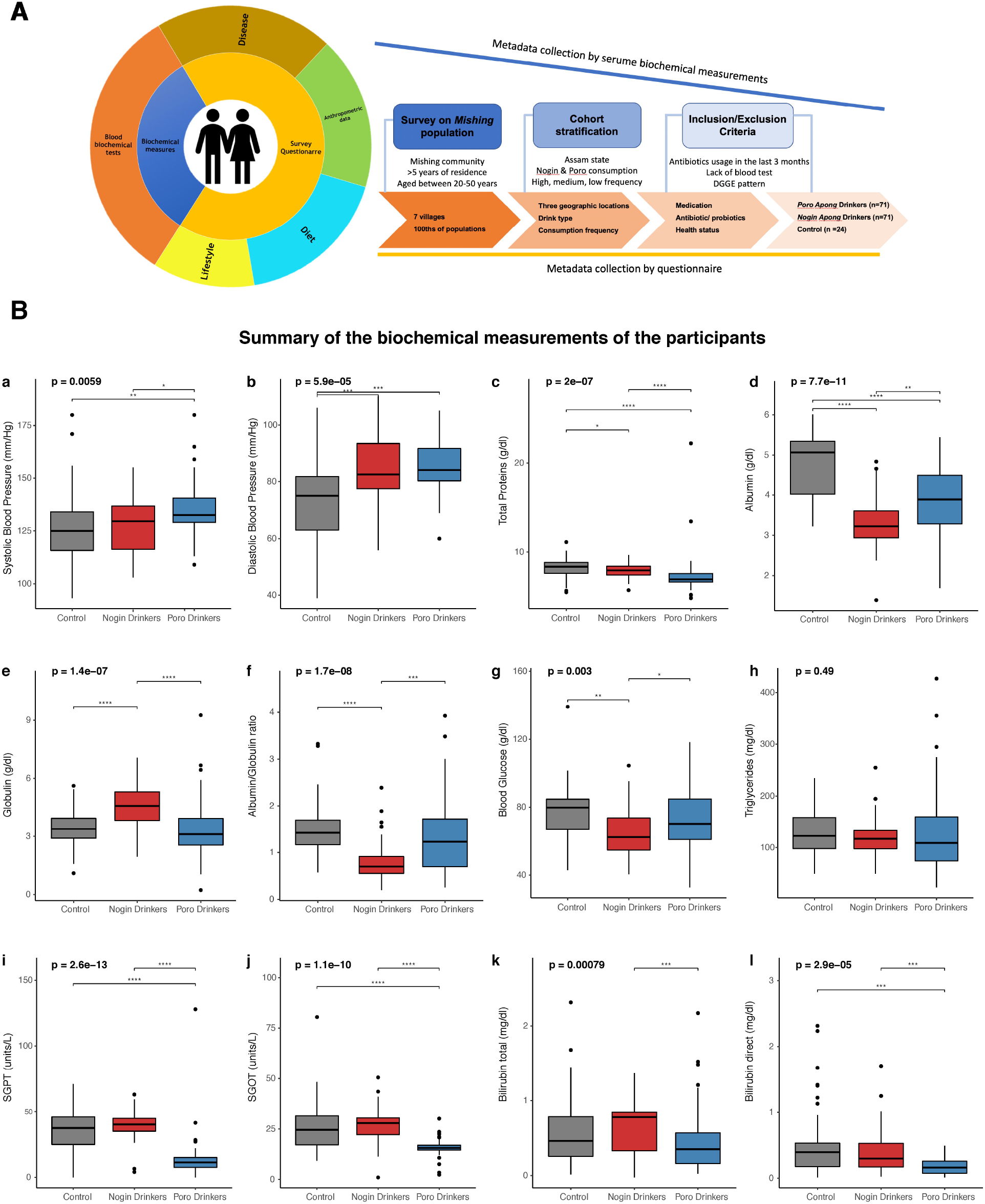
**A)** Summary of the cohort. **B)** A comparison of anthropological factors and serum biochemical markers among *Apong* drinkers and non-drinkers (a) Systolic Blood pressure, (b) Diastolic Blood pressure, (c) Total protein (d) Albumin, (e) Globulin, (f) Albumin/Globulin ratio, (g) Blood Glucose, (h) Triglycerides, (i) serum glutamic-pyruvic transaminase SGPT, (j) Serum glutamic-oxaloacetic transaminase (SGOT), (k) Bilirubin total and, (l) Bilirubin direct. The probability of significance is denoted by *’ s, where **** depicts significance level of 0.0001, *** depicts 0.001, ** depicts 0.01, * depicts 0.05. Only significant p-values are indicated.

### The gut microbiota composition varies between non-drinkers and *Apong* drinkers

We compared the gut microbial composition of *Apong* drinkers and non-drinkers. We found significant differences between *Nogin* and *Poro* drinkers and drinkers and non-drinkers **(Figure 2A)**. Overall, the participants were dominated by *Bacteroidota* and *Firmicutes*, with smaller proportions of *Proteobacteria*. However, while *Bacteriodata* made up a significant proportion of the gut bacterial community in *Nogin* drinkers and non-drinkers, it made up a smaller proportion in *Poro* drinkers. Notably, *Actinobacteria* was only detected in non-drinkers. Next, we identified taxa at the family level that significantly differed in abundance among the *Apong* drinkers and non-drinkers **(Figure 2B)**. *Apong* drinkers had higher colonization by Enterobacteriaceae and Ruminococcaceae compared to non-drinkers. On the other hand, *Poro* drinkers had significantly less Prevotellaceae than *Nogin* drinkers and non-drinkers.

**Figure 2.**
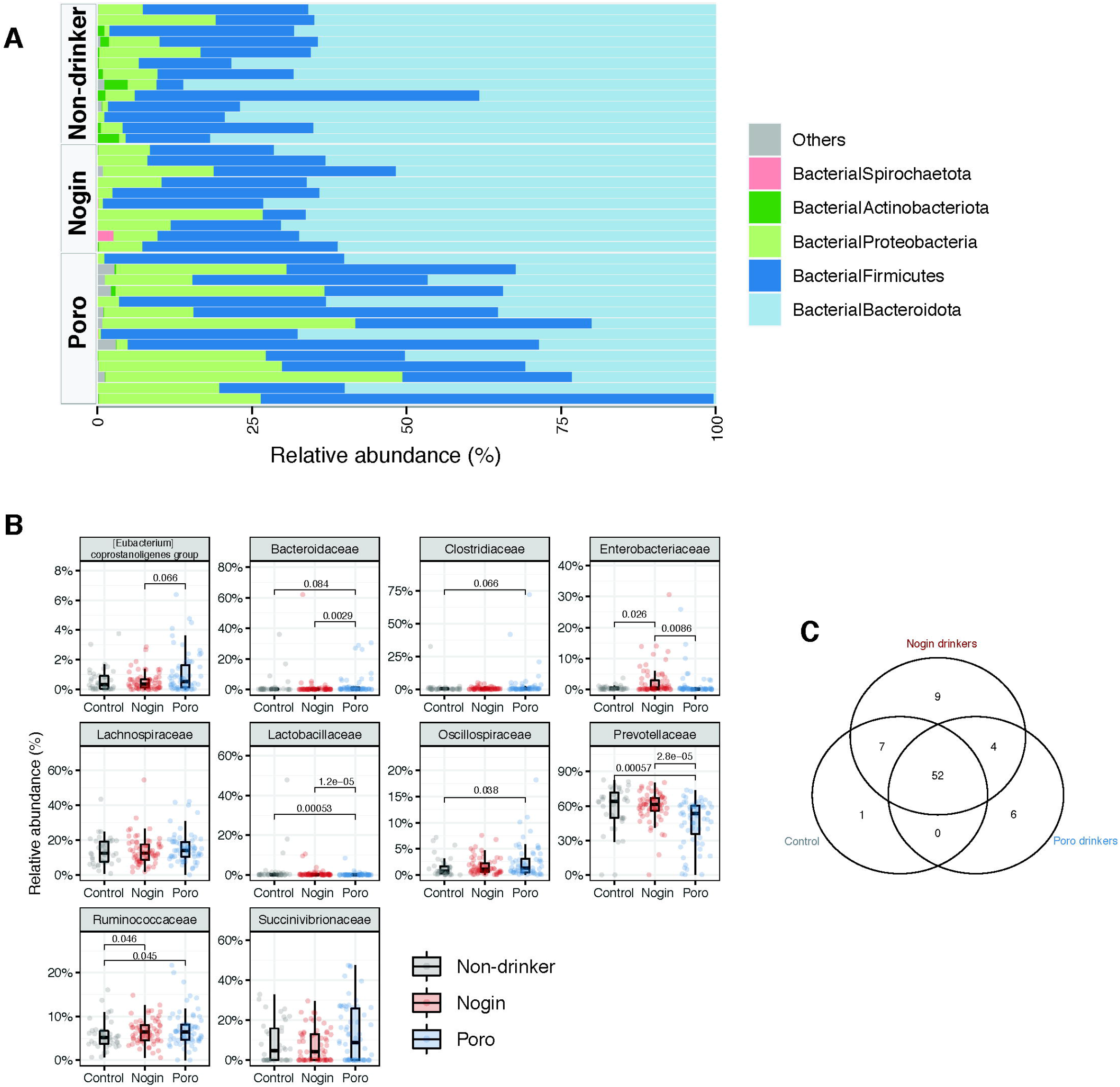
**A)** Relative abundance of microbial taxa at phylum level in *Apong* drinkers and non-drinkers. **B)** Top ten microbial families that are differentially abundant among *Apong* drinkers and non-drinkers. Only significant p-values are indicated. **C)** Venn diagram of microbial features (ASVs) shared between different drink type groups or special to each group.

We also identified microbes that were specific to *Apong* drinkers **(Figure 2C)**. To do this, we identified and compared ASVs that were detected in at least 50% of all participants at a minimum abundance of 0.1% in *Apong* drinkers and non-drinkers. Most of the ASVs were shared (n=52) among all participants and were part of the “core” microbiome. However, certain ASVs were only detected in *Nogin* (n=9) or *Poro* drinkers (n=6). Four of the *Poro*-specific ASVs were from the Lachnospiraceae family, while the others were from the Succinivibrionaceae family and an unknown family of Bacteroidales. The majority of the *Nogin*-specific ASVs were from the Prevotellaceae family (n=7), with the remaining ASVs coming from the *Ruminococcaceae* and *Enterobacteriaceae* families. The only ASV that was detected in non-drinkers but not in *Apong* drinkers was from the *Prevotellaceae* family.

### Gut microbial diversity in *Apong* drinkers is higher than in non-drinkers, and highfrequency Nogin drinkers have lower gut microbial diversity than other Nogin drinkers

We estimated the gut microbial diversity in *Apong* drinkers and non-drinkers using three different diversity metrics **(Figure 3A)**. Gut microbial diversity was significantly higher in the *Apong* drinkers than the non-drinkers.

**Figure 3.**
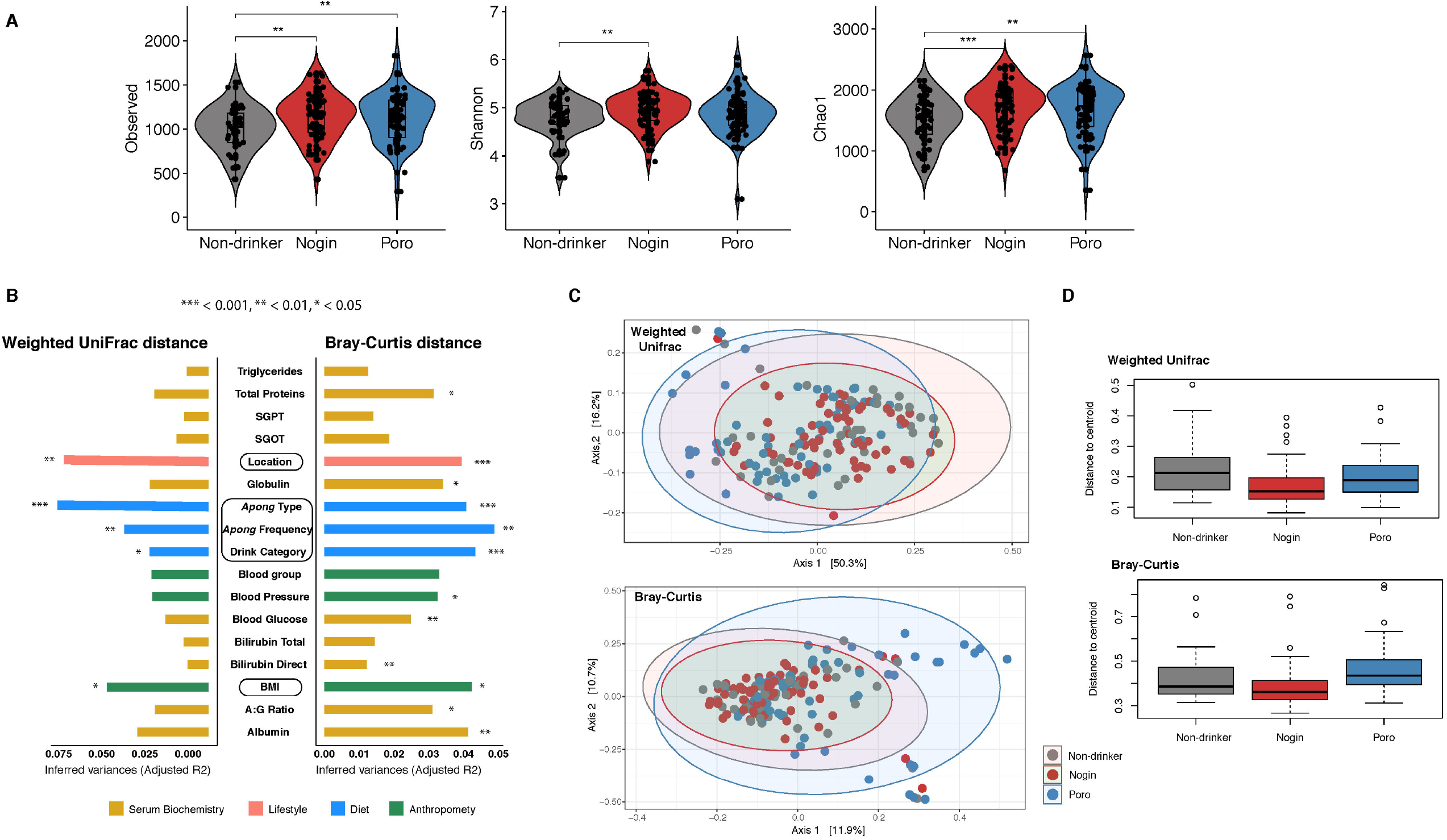
**A)** Microbial diversity of *Apong* drinkers and non-drinkers as estimated by ASV richness and Shannon and Chao1 indices. **B)** The inferred variance (adjusted R2) explained by each identified covariate as determined by PERMANOVA, calculated based on weighted UniFrac and Bray–Curtis dissimilarities. Statistically significant covariates with an adjusted *p*lJ<LJ0.05 using the Benjamini–Hochberg (BH) method are shown. **C)** PCoA of the weighted UniFrac and BrayCurtis distances of the gut bacterial composition of *Apong* drinkers and non-drinkers. **D)** Dispersion (i.e. distance to centroid of the groups) of each drink type group in the PCoA plots.

We divided the participants into three categories based on their *Apong* consumption frequency: high (HD), medium (MD), and low (LD). *Poro* consumption frequency did not have an effect on microbial diversity **(Supplementary Figure 1)**. However, among Nogin drinkers, highfrequency drinkers had significantly lower gut microbial diversity.

### *Apong* consumption and frequency has a significant effect in gut microbial composition

We investigated microbial composition between participants by computing Bray-Curtis and weighted UniFrac beta-diversity **(Figure 3)**. The distance matrices revealed small but significant differences among the non-drinkers, *Nogin* drinkers, and *Poro* drinkers, as well as among subgroups of *Apong* drinkers based on frequency of consumption **(Figure 3A)**.

Although location also had a significant effect on microbial composition, *Apong* usage had a larger overall effect. **(Figure 3A)**. When blocking “location” in the perMANOVA, drink type (non-drinker vs. *Nogin*, or *Poro* drinker) still explains a significant portion of the variation in microbial composition (Bray-Curtis: R^2^= 0.034; p= 0.003 & Unifrac: R^2^= 0.035; p= 0.001). Despite inhabiting the same location (Majuli), the gut microbial composition of non-drinkers (n=24) and *Nogin* drinkers (n=18) formed two different clusters of PCoAs (R^2^= 4.79; p=0.005) **(Supplementary Figure 2)**.

Notably, blood serum biochemistry markers, such as total protein, albumin and globulin levels, and blood glucose and pressure, explained significant variation in the presence or absence of certain microbes, but not in the phylogenetic diversity of gut microbial composition.

### *Nogin* drinkers have a more homogenous gut microbial community than *Poro* drinkers and non-drinkers

The gut microbial community of *Nogin* drinkers clustered together in the principal coordinate analysis (PCoA) based on both weighted UniFrac and Bray-Curtis distances **(Figure 3C)**. However, non-drinkers and *Poro* drinkers had more heterogeneous microbial communities than *Nogin* drinkers. This was also demonstrated by the higher distance to centroid (Multivariate homogeneity of groups dispersions) **(Figure 3D)**, where *Nogin* drinkers had the lowest distance to centroid, so highest homogeneity whereas *Poro* drinkers had a high distance, so more heterogeneity in the weighted UniFrac distance (considers phylogenetic relatedness).

### Fecal microbial metabolite profiles are different between the *Apong* drinkers and nondrinkers

To study the metabolic activity in the gut ecosystem of the cohorts, an untargeted metabolite profiling with GC-MS analysis was performed. Metabolites of microbial origins were identified using the human metabolome database (HMDB) for subsequent analysis. We extracted a total of 384 metabolites which comprises mainly amino acids, bile acids, fatty acids, indoles, and saccharides. *Apong* consumption led to depletion of certain metabolites, such as acetamid, benzestrol, butanedioic acid, and cyclopropanecarboxylic acid, while *Poro* drinkers had higher levels of undecanoic acid than non-drinkers and *Nogin* drinkers **(Figure 4)**.

**Figure 4.**
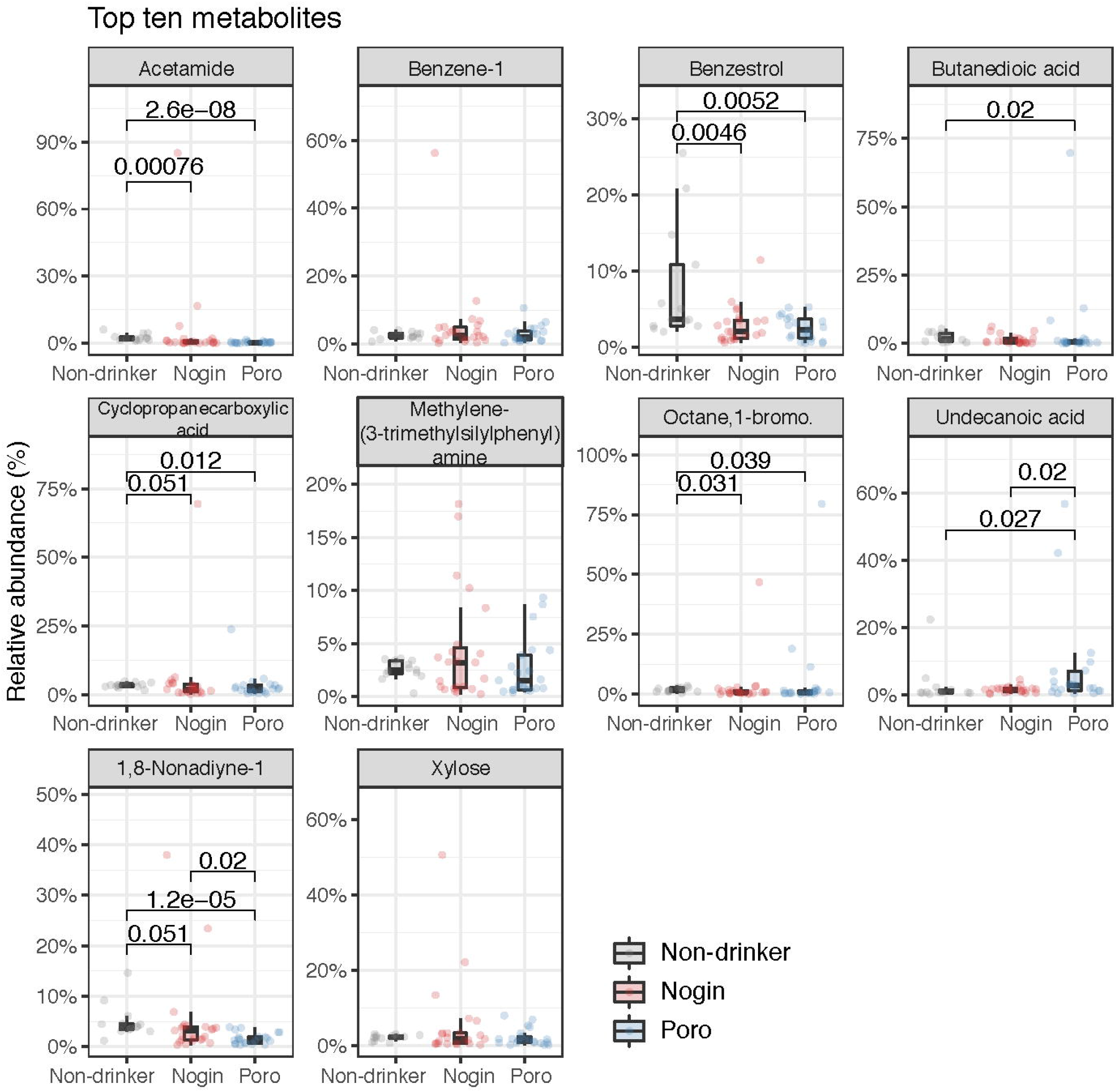
Top ten fecal metabolites that are differentially abundant among *Apong* drinkers and non-drinkers. Only significant p-values are indicated.

We agglomerated gut microbes at family level and correlated these families to fecal metabolites using the mbOmic package in R^17^. We found 113 significant correlations (adjusted p-values < 0.05) between microbial families and fecal metabolites. Only 23 of these correlations had a rho value higher than 0.70. Top ten correlations are listed in **Figure 5A**, and the majority of these correlations were with Clostridia or unknown Bacteroidota. All the correlations are listed in **Supplementary Table 1**.

**Figure 5.**
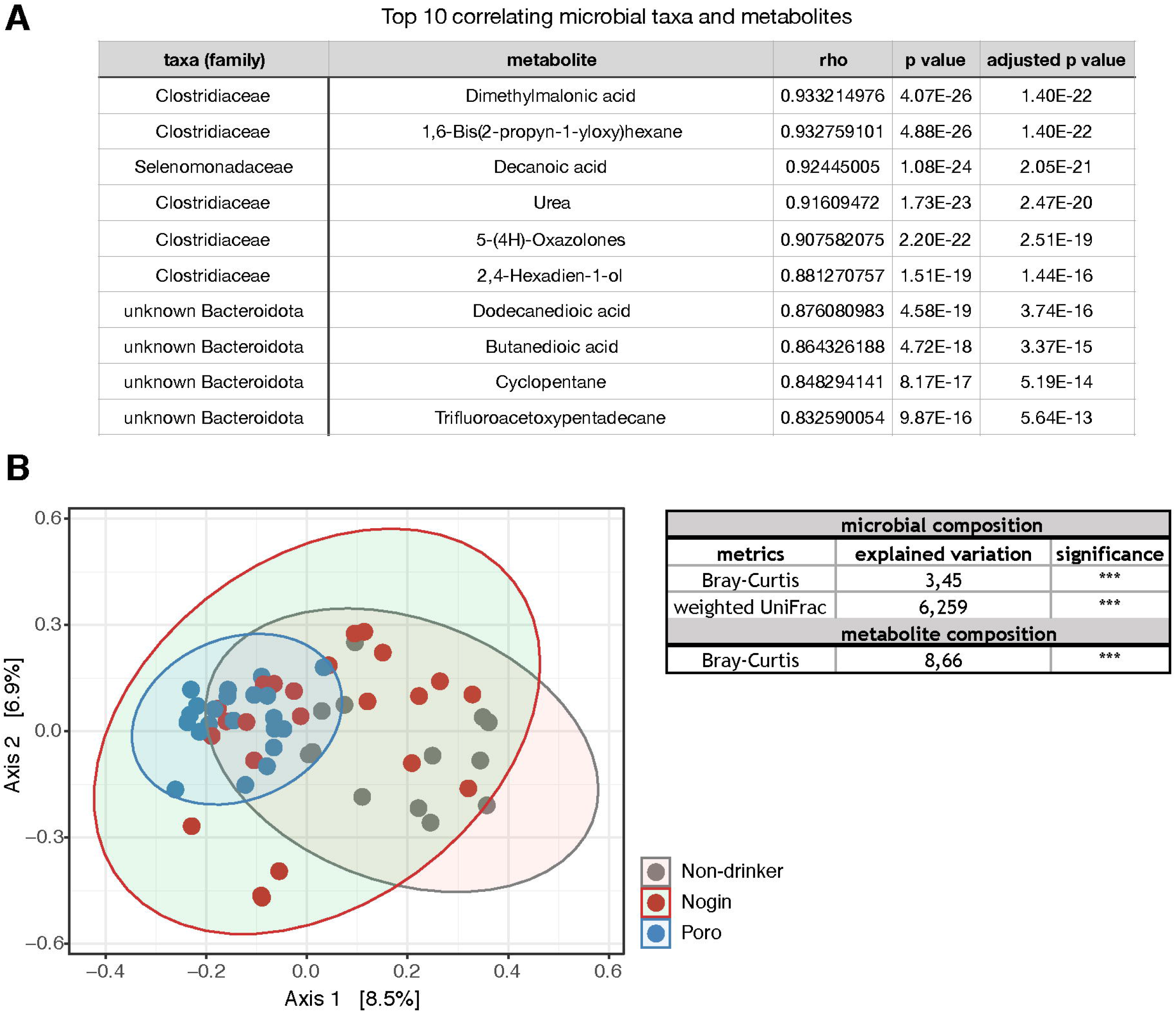
**A)** Top ten highest correlations between gut microbial taxa at family level and fecal metabolites. **B)** PCoA of the Bray-Curtis distances of the fecal metabolites of *Apong* drinkers and non-drinkers. The table shows explained variance by drink type (*Nogo, Poro*, and non-drinker) for microbial and fecal metabolites composition.

Although having relatively heterogeneous gut microbial composition among participants **(Figure 3C-D)**, *Poro* drinkers had a highly uniform composition of fecal metabolites **(Figure 5B)**. Drink type (non-drinker, *Nogin* drinker, or *Poro* drinker) had even a larger effect on metabolite composition than on gut microbiome composition **(Figure 5C)**.

Taken together, these results show that *Apong* drinkers had distinct blood serum markers, gut microbial composition, and fecal metabolites compared to non-drinkers.

### *Apong* consumers had lower levels of iso-valeric acid among their fecal SCFAs compared to non-drinkers

The gut microbiota helps to break down undigested food through fermentation, producing short-chain fatty acids (SCFAs) as a result^18^. The SCFAs are important for gut homeostasis and health and their levels are affected by diet. Although BCFAs (branched-chain fatty acids) may be important in the gut and could potentially serve as markers of gut microbial metabolism, they have received less attention than the major SCFAs^19^.

We measured the levels of three major SCFAs and one BCFA (acetic acid, butyric acid, propionic acid, and iso-valeric acid) in fecal samples from both *Apong* drinkers and non-drinkers using HPLC. Propionic acid was the most abundant SCFA in both groups, while butyric acid was the least **(Figure 6)**. Some volunteers had very high levels of acetic acid, but it was not correlated with any other data **(Supplementary Figure 3)**. Isovaleric acid, a BCFA and considered to be harmful to the colon epithelium^20^, was significantly lower in *Apong* drinkers compared to non-drinkers, but there was no significant difference for the other SCFAs **(Figure 6)**.

**Figure 6.**
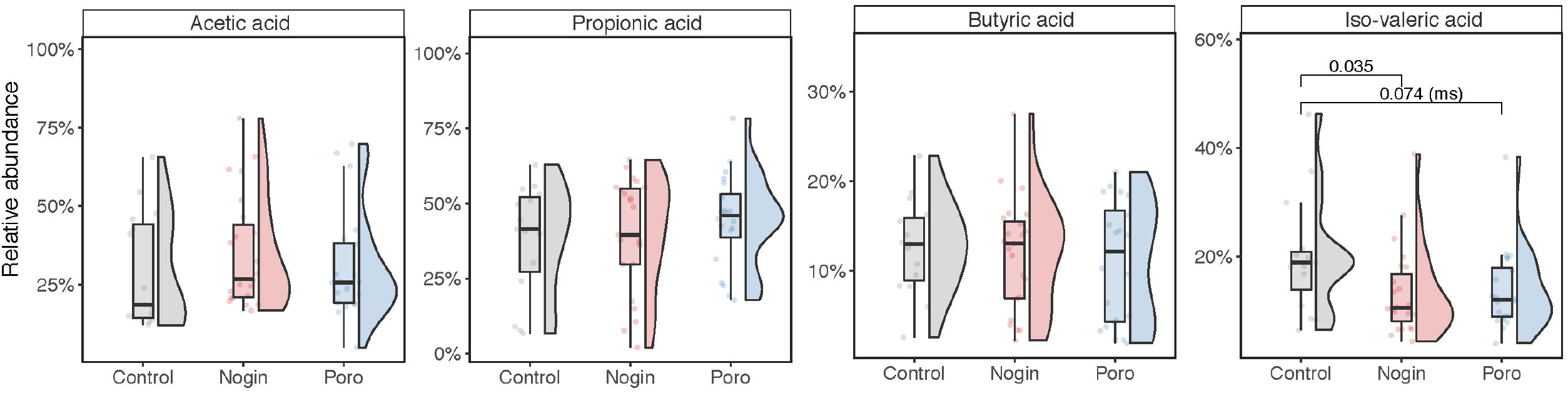
Composition of the four short chain fatty acids (SCFAs) in the fecal samples of *Apong* drinkers and non-drinkers. Only significant p-values are indicated. _“_ms” denotes marginal significance.

## Discussion

The composition of the gut microbiome is influenced by various factors, including diet and life-style^4, 21–23^, but the impact of a single component of diet within a population of the same ethnicity on the gut microbiome has not been studied before. This study investigates the effect of two types of a traditional, rice-based fermented alcoholic beverage (*Apong*) on the gut microbiome and health of the *Mishing* community in India. All volunteers in the study were of the same ethnicity, healthy, and had similar dietary habits, except for being an *Apong* drinker or not.

Our study showed that the composition of the gut microbiome is affected by the choice of *Apong*. Despite the frequent consumption of an alcoholic beverage, the volunteers in the study had normal BMI and healthy vital organ function. Previous research has shown that both varieties of *Apong* contain mild alcohol (~10%) and have a high content of phenolics^14^.

This study found that *Apong* consumers had a more diverse gut microbiome compared to the non-drinkers, which had a stable community with fewer ASVs. This suggests that a single dietary factor alone can significantly impact the gut microbiome of a community, consistent with previous research on the gut microbiome of children in Burkina Faso^24^. Our study also observed a strong association between variation in microbial composition and blood glucose levels and blood pressure, which has not been reported in previous studies. Further research is needed to understand the causal relationship between these factors and gut bacterial diversity, which may allow for the development of microbiome-based biomarkers for predicting lifestyle diseases.

In this study, the gut microbiome of the *Mishing* community was found to be dominated by the *Prevotellaceae* family, a signature of the Indian population^25–30^, which has been previously associated with a vegetarian or carbohydrate-rich diet^7^. However, *Poro* drinkers had lower levels of *Prevotellaceae* than non-drinkers and *Nogin* drinkers, even though their gut microbiomes were colonized to a high extent by *Prevotellaceae*.

Lastly, we found that the gut microbiome of the *Mishing* population was colonized to a high extent by *Succinivibrio*, a bacterium not previously reported in the Indian population^25–27^. This bacterium is commonly found among hunter-gatherers and foragers^31, 32^. The presence of *Succinivibrio* in the microbiome was previously reported in the *Hadza* hunter gatherers and traditional Peruvian populations^31,33^. We speculate that the high abundance of *Succinivibrio* in the *Mishing* population may be due to co-habitation with domesticated animals. The co-presence of butyrate producers and other essential gut bacteria along with expected levels of SCFAs but lower levels of BCFA and blood serum measurements suggests that *Apong* does not have a detrimental effect on the structure and function of the gut microbiome.

## Conclusion

In conclusion, this study found that the choice of *Apong*, a traditional rice-based fermented alcoholic beverage, significantly impacts the gut microbiome composition and blood serum markers in the *Mishing* community in India. The gut microbiomes of *Apong* consumers were more diverse than those of non-drinkers, and *Poro* drinkers had lower levels of Prevotellaceae than non-drinkers and *Nogin* drinkers. The gut microbiome of the *Mishing* community was also colonized by *Succinivibrio*, a bacterium not previously reported in the Indian population. The differences in gut metabolites between *Apong* drinkers and non-drinkers were even greater than those in the gut microbiome. These findings suggest that a single dietary factor can significantly impact the gut of a population and highlight the need for further research on the causal relationship between these factors and gut bacterial diversity for the development of microbiome-based biomarkers for predicting lifestyle diseases.

## Materials and methods

### Materials

All the reagents, media and chemicals used in this study were of analytical grade. RNA-later solution (Cat. No RM49049), short chain fatty acids standards viz. Acetic acid (Cat. No. 5438080100), Butyric acid (Cat. No 19215), Isovaleric acid (Cat. No 78651), and Propionic acid (Cat No. 94425) were procured from Sigma-Aldrich, USA. QiAmp DNA stool mini kit (Cat. No: 19590) was procured from Qiagen Inc, Germany. Blood collection vials K3-EDTA vial (Cat. No 368860) and clot vials (Cat. No 368975) were procured from BD Diagnostics, Oxford, UK. Blood serum biomarkers kits were procured from CCS ® Coral Clinical Systems, Tulip Diagnostics (P) Ltd., Goa, India.

### Ethics statement

This study was approved by the Ethics Committee of the Institute of Advanced Study in Science Technology (IASST), Guwahati, India (Approval number: IEC(HS)/IASST/1082/2014-15/6) and was conducted following the guidelines and regulations. Written informed consents were taken from the volunteers along with some standard questionnaires.

### Study sites, volunteers, and sample collection

In this study, potential volunteers were identified through an electoral database and surveyed in locations dominated by the *Mishing* community. The control group consisted of individuals who do not consume *Apong* but follow the same dietary pattern, while the experimental group included 166 individuals from the *Mishing* community and 24 individuals from a Vaishnavite satra with slightly different dietary habits. A bi-lingual survey questionnaire was used to collect information on dietary patterns, age, sex, medical history, family lineage, and demographics. Inclusion criteria were age (18-50 years) and ethnicity, while exclusion criteria were the use of antibiotics, health supplements and other drugs, consumption of any other liquor except *Apong*, and medical history. The *Mishing* population was stratified based on the amount of rice beer consumed per week, with non-drinkers classified as those who did not consume *Apong*, low-drinkers as those who consumed less than 250 ml, medium-drinkers as those who consumed 250-500 ml, and high-drinkers as those who consumed more than 500 ml per day. The control group included monks from *Vaishnavite* satras, who avoided alcoholic beverages and red meat but consumed fish as part of their regular diet.

Fecal and blood samples were collected from the Mishing population living in the Telam, Dhemaji, and Majuli districts of Assam. A customized kit was used to collect fecal samples, with one container, including 2 ml of RNA later solution, used for DNA extraction and another for metabolomics studies. Three ml of blood was withdrawn from individuals by a phlebotomist. Blood was collected in a K3-EDTA vial and clot vials to separate plasma and serum, respectively. All the samples were immediately frozen after collection and were transported to the laboratory in frozen condition. Fecal and blood samples were stored at −80 °C, until processed. In addition to the samples and questionnaires, anthropometric data such as height, weight, BMI, and blood group were also collected from the volunteers.

### Analysis of biochemical parameters of plasma and serum

The plasma samples were analyzed using standard biochemical assay kits (CCS ® Coral Clinical Systems, Tulip Diagnostics (P) Ltd., Goa, India), following the manufacturer’ s instructions. Serum glucose, HDL cholesterol, albumin, globulin, total protein, and liver function tests, for example, serum glutamic oxaloacetic transaminase (SGOT), serum glutamic pyruvic transaminase (SGPT), and alkaline phosphatase (ALP), direct and total bilirubin contents were determined.

### DNA extraction from faecal sample

DNA extraction was performed within 72 h of faecal sample collection. QiAmp DNA stool mini kit (Cat. No: 19590 Qiagen Inc, Germany) was used for the extraction of metagenomic DNA from the faecal samples following the manufacturer’ s instruction. Briefly, a 400 µl of faecal sample was mixed to 1400 µl of ASL buffer (provided with the kit) and incubated at 90 °C for 10 min. The supernatant was collected after brief centrifugation at 13,000 rpm for 2 min, to which Inhibitex tablet and ProteinaseK (supplied with the kit) were added. After a short incubation of 10 min at 70 °C, the mixture was centrifuged at 13,000 rpm for 3 min. The supernatant was collected in a filter with a silica column. The column was washed twice with a wash buffer (provided with the kit) and the bound DNA was eluted with a preheated elution buffer (supplied with the kit). The amount of dsDNA was quantified using a dsDNA estimation kit with a Fluorometer (Quantiflour, Promega, Madison, USA).

### Library preparation, 16S rDNA amplicon sequencing and analyses

The V3-V4 region of the 16S rDNA was amplified using the primer pairs 341F & 805R^34,41^. The indexing and library preparation of the amplified DNA fragments were carried out using Nextera XT library preparation and indexing kits according to the Illumina MiSeq protocol^35^. DNA fragments were multiplexed and subjected to 2 × 300 bp paired-end sequencing in an Illumina MiSeq machine with the sequencing service provider (Macrogen Inc., Seoul, Republic of Korea).

### Bioinformatics analyses of the amplicon dataset

The paired-end reads generated from Illumina sequencing were processed using the LotuS2 pipeline^36^. Reads having less than 170 bases in length were filtered out from the analysis. In LotuS2, DADA2 algorithm^37^ was used to cluster sequences into amplicon sequence variants (ASVs). Using the options for LULU (-lulu)^38,45^ and UNCROSS2 (-xtalk), sequence clusters were curated and refined. ASVs were aligned with Lambda^39^ to SILVA 138.1^40^, to obtain taxonomic assignments for ASVs using the LotuS2 LCA algorithm. Otherwise, default options in LotuS2 were used. As a result, 28,083,099 total reads were clustered into 7550 ASVs in the final matrix, summing to 14,893,374 reads after quality filtering and removal of contaminants^41^.

The processed samples were further analysed with phyloseq package^42^ in R (version 3.6.1). Samples were rarefied to an even depth using the rtk tool^43^ for diversity analysis. For estimation and calculation of diversity indices, vegan^44^ package was used.

### Metabolomics analyses of fecal samples

The following techniques were used for the analyses of fecal samples.

### Gas chromatography-mass spectroscopy (GC-MS) analysis

Untargeted fecal metabolite profiles were determined by GC-MS analysis. A 40 mg of lyophilized fecal sample was extracted with 1 ml of HPLC grade methanol (Merck, Mumbai, India) and kept in a shaker overnight at 1200 rpm. The sample was centrifuged at 10000 rpm for 10 min at 10°C. The extract was then dried at room temperature using a vacuum desiccator and resuspended in a mixture of 40 µl of pyridine and 20 mg/ml of methoxyamine hydrochloride. After a brief vortex, the solution was incubated at 30 °C for 90 min and then derivatized with 20 µl of N-methyl-N-trimethylsilyltrifluoroacetamide with 1% trimethylchlorosilane (Merck, USA) at 70 °C for 30 min^54^. The sample was then centrifuged at 3000 rpm for 5 min and used for GC-MS analysis.

Samples were run in a Shimadzu GC 2010Plus-triple quadrupole (TP-8030) system fitted with an EB-5MS column (length: 30 m, thickness: 0.25 μm, ID: 0.25 mm). A 1 µl of the sample was injected in split less mode at 300°C using helium as carrier gas at a 1 ml/min flow rate. The oven program was set at 70°C initially, ramped at 1°C/min for 5 min up to 75°C. Subsequently, it was increased at 10°C/min up to 150 °C, and held for 5 min, followed by increasing at the same rate up to 300 °C and held for 5 min. The mass spectrometer was operated at a continuous scan from 45 to 600 m/z in the electron ionization (EI) mode at 70ev with 200 °C as the source temperature. Peak identification was performed using the National Institute of Standards and Technology library, USA, by matching the mass spectra.

### Quantification of faecal short-chain fatty acids (SCFAs) by RP-HPLC analysis

Faecal samples were lyophilized, and 100 mg of each sample was dissolved in a solvent prepared using acetonitrile (Cat. No 271004 Merck Millipore, Germany) and 10-mM KH_2_PO_4_ (pH 2.4) (Cat. No P5379, Sigma Aldrich, USA) in 1:1 ratio. The solutions were then vortexed vigorously for 5 min. ensuring proper mixture, centrifuged at 4000 rpm for 5 min. The supernatant was collected and filtered through a 0.22 µm syringe filter for downstream analysis.

Quantification of the organic acids was carried out in an analytical HPLC instrument (Waters, USA) with 5 µm ODS2 (4.6 × 250 mm, Waters SPHERISORB) reversed- phase C18 analytical column. Two HPLC grade solvents (solvent A was 10-mM KH_2_PO_4_, pH 2.4 with phosphoric acid, while solvent B was 100% acetonitrile) were used in a gradient system with a flow rate of 1.5 ml/min. The absorbance of the eluted compound was monitored at 210 nm by the PDA detector.

### Statistical analyses

All the statistical tests were performed in the R platform by using base functions and calling specialized packages such as phyloseq^42^, vegan^44^, microeco^45^, microbiome^46^, microbiomeutilities^10^, mbOmic^17^, rstatitix^51^. Comparisons among the anthropometric measures and serum biochemical markers were carried out using the Kruskal-Wallis test. To compare the microbial diversity among *Apong* drinkers and non-drinkers, the samples were first rarefied to an equal depth. Then, the alpha diversity indices were calculated for each sample, including the number of unique features (richness), the Shannon diversity, and Chao1 metric. To determine if there were significant differences in microbial diversity among *Apong* drinkers and non-drinkers, a Kruskal-Wallis test was performed. Before calculating the beta distance (Bray-Curtis and weighted UniFrac), the table was normalized to relative abundances. “adonis” and “mantel” functions in the vegan package were used to run PERMANOVA (Permutational Multivariate Analysis of Variance) and Mantel tests to calculate metadata variables explaining variation based on beta diversity distances. _“_betadisper” function in the vegan package was used to estimate the homogeneity of drink types groups in the PCoAs. The “corr” function from the mbOmic package was used to calculate Pearson correlations between the metabolomics data and microbial taxa at the family level and adjusted p values for multiple comparisons were used.

## Acknowledgement

The authors acknowledge the Institutional Level Biotech Hub (DBT, Govt. of India) and Central instrumentation facility of IASST for providing the facilities. This research was funded by Department of Biotechnology (DBT) under Unit of Excellence Project (BT/550/NE/U-Excel/2014) and SC/ST community development programme in IASST (SEED/TITE/2019/103) funded by DST, Govt. of India. SD is thankful to IASST for providing financial support to carry out research work. The authors also acknowledge Mr. Pinku Rajbongshi for assisting in collection of blood samples. We are thankful to Mr. Dhrubajyoti Regon, Mr. James Doley, Mr. Deepak Mili and Mr. Samujjal Saikia for assistance in recruitment of volunteers and sample collection. EO, FH were supported by European Research Council H2020 StG (erc-stg-948219, EPYC). CF, RA, EO, FH were supported by the Biotechnology and Biological Sciences Research Council (BBSRC) Institute Strategic Programme Gut Microbes and Health BB/r012490/1 and its constituent project BBS/e/F/000Pr10355, Core Capability Grant BB/CCG1720/1 and the work delivered via the Scientific Computing group, as well as support for the physical HPC infrastructure and data centre delivered via the NBI Computing infrastructure for Science (CiS) group.

## Availability of data

Sequencing data are available on the NCBI SRA server under the BioProject ID: PRJNA906264

## Figures legends

**Supplementary Figure 1:** Microbial diversity of *Apong* drinkers estimated by ASV richness: _“_low-drinkers” (less than 250 ml per day), _“_medium-drinkers” (250-500 ml per day), and _“_highdrinkers” (more than 500 ml per day).

**Supplementary Figure 2:** PCoA of the weighted UniFrac and Bray-Curtis distances of the gut bacterial composition of *Apong* drinkers and non-drinkers where colors depict *Nogin, Poro*, and non-drinkers and shapes depict different locations.

**Supplementary Figure 3:** Composition of the four short chain fatty acids (SCFAs) in the fecal samples of the participants

**Supplementary Table 1:** Correlations between gut microbial taxa at family level and fecal metabolites.

