## Supplementary figures and images for "A single dietary factor, daily consumption of a fermented beverage, can modulate the gut microbiome within the same ethnic community"

### Supplemental Figure 1

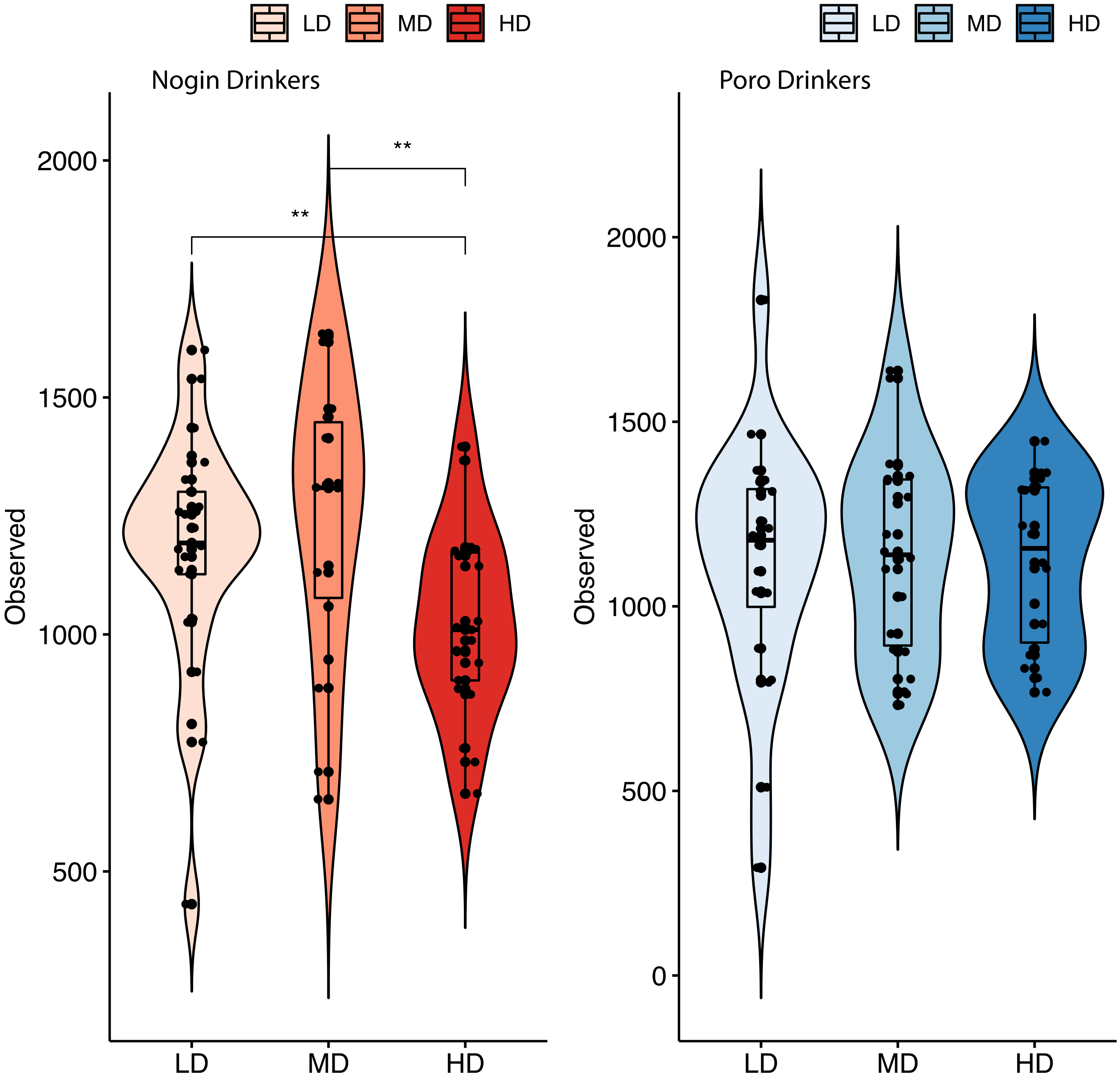

### Supplemental Figure 2

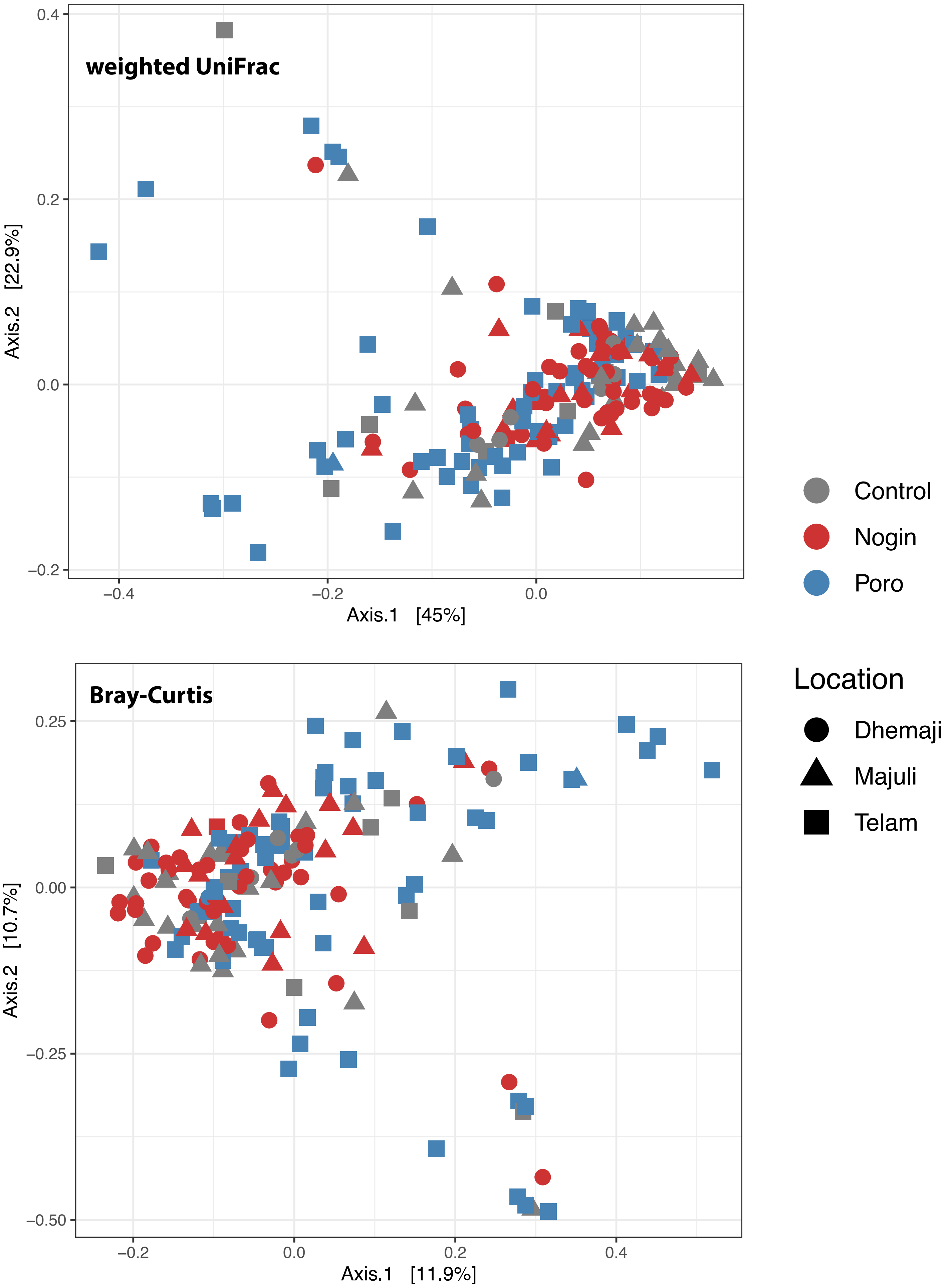

### Supplemental Figure 3

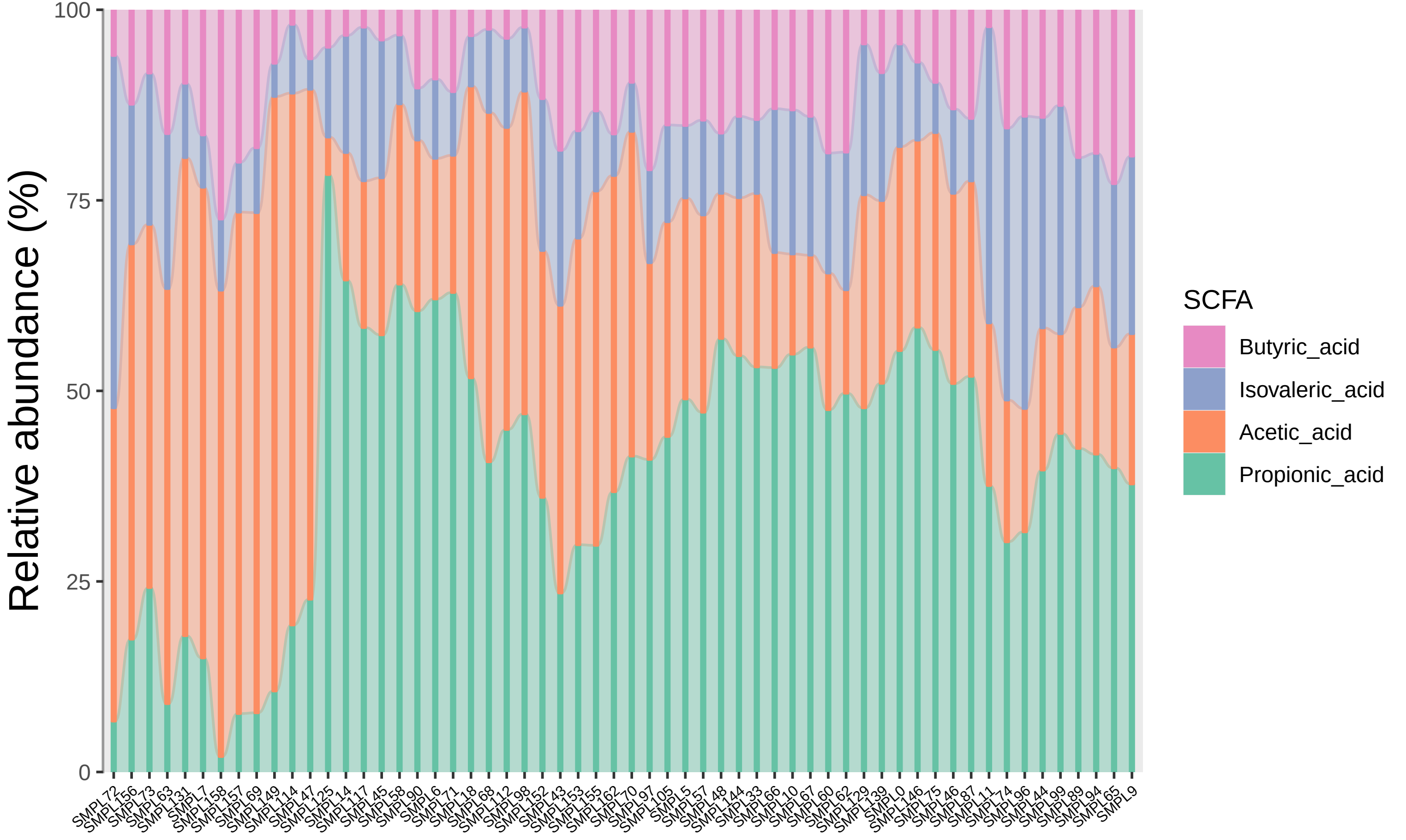
